# Unveiling Frailty: Comprehensive and Sex-Specific Characterization in Prematurely Aging PolgA Mice

**DOI:** 10.1101/2024.02.15.580562

**Authors:** Dilara Yılmaz, Amit Singh, Esther Wehrle, Gisela A. Kuhn, Neashan Mathavan, Ralph Müller

## Abstract

Frailty, a geriatric syndrome, is assessed using the frailty phenotype (FP) and frailty index (FI). While these approaches have been applied to aging mice, their effectiveness in prematurely aging mouse models such as PolgA^D257A/D257A^ (PolgA) has not been completely explored. We demonstrated that frailty became evident in PolgA mice around 40 weeks, validated through body weight loss, reduced walking speed, decreased physical activity, and weaker grip strength. Moreover, we also identified sex differences in these mice with females exhibiting higher frailty compared to males. Frailty prevalence in PolgA mice at 40 weeks parallels that observed in naturally aging mice at 27 months and aging humans at 65-70 years. These findings contribute to understanding frailty onset and sex-specific patterns, emphasizing the significance of the PolgA mouse model in investigating aging and related disorders.

## Introduction

Frailty is a complex geriatric syndrome that is characterized by a diminished ability to respond to stressors and maintain homeostasis (Milte and Crotty, 2014). Two widely used approaches to evaluate frailty are the frailty phenotype (FP) and the frailty index (FI) (Fried et al., 2001, Rockwood et al., 2005). The FP assesses individuals based on the presence or absence of specific criteria, which are: unintentional weight loss, slow walking speed, self-reported exhaustion, weakness, and low physical activity. Individuals, meeting three or more of these criteria are classified as frail [2]. On the other hand, the FI is based on the accumulation of health deficits, where the total number of deficits is divided by the number of deficits assessed, providing a quantitative measure of frailty (Rockwood et al., 2005). Both the FP and FI are valuable approaches for identifying frailty in individuals, enabling a better understanding of its impact on health outcomes and aging research.

With the intention of investigating frailty, mouse models that accurately replicate the complex nature of frailty in humans have been developed (Yilmaz et al., 2023). Several research groups established frailty assessment tools designated for mice based on human criteria (FP or FI)

(Baumann et al., 2016, Baumann et al., 2020, Liu et al., 2014, Parks et al., 2012). Slowness (rotarod), weakness (grip strength), physical activity (rotarod), and endurance (combined rotarod and grip strength) were assessed to develop frailty in mice by adopting the FP approach for humans (Liu et al., 2014). Similarly, frailty was defined in mice by using the clinical human FI approach, which involved 31 parameters, including basal metabolism, body composition, and activity levels in evaluating signs of discomfort (Whitehead et al., 2014) and the modified versions of these methods were used by other groups to assess frailty (Baumann et al., 2018, Baumann et al., 2019, Baumann et al., 2020, Gomez-Cabrera et al., 2017, Kwak et al., 2020, Sukoff Rizzo et al., 2018). However, the methods employed for assessing frailty in mice have primarily been applied to C57BL/6 (naturally aging) mice aged 17 months and older (von Zglinicki et al., 2016). Therefore, it remains unclear whether these methods are equally effective in identifying frailty in prematurely aging mice.

In this regard, prematurely aging mouse models such as PolgA^D257A/D257A^ (PolgA) offer numerous advantages over naturally aging mice (Yilmaz et al., 2023). Notably, PolgA mice demonstrate the premature manifestation of age-related phenotypes within a short time frame (Trifunovic et al., 2004). Around just 40 weeks of age, these mice exhibit multiple hallmarks of aging, including kyphosis, hearing impairment, progressive weight loss, osteosarcopenia, and osteoporosis (Kujoth et al., 2006, Trifunovic et al., 2004, Scheuren et al., 2020a). In addition, age-related changes in these mice were evaluated in females using the clinical mouse FI method, exhibiting higher FI scores in PolgA mice compared to their age-matched wild-type littermates (WT) (Scheuren et al., 2020a). Although the PolgA mouse model holds promise for investigating frailty and age-related musculoskeletal disorders, there are certain limitations in its frailty scoring. Despite being a non-invasive method, FI scoring, in general, lacks assessments such as walking speed and endurance to measure the functional abilities of mice. Moreover, the sex-dependent component of frailty has not been addressed in these mice. Therefore, our first objective is to identify the frailty phenotype in prematurely aging PolgA mice around 40 weeks by evaluating weight loss, grip strength (weakness), walking speed (slowness), and home cage walking distance (physical activity) in comparison to their age-matched WT littermates. Our second objective is to show the sex-dependent differences in frailty in PolgA mice.

## Results

When considering age-related changes in the PolgA mice, there were notable differences compared to WT mice for both sexes (Figure 1). PolgA males weighed 13% less compared to their WT counterparts (p<0.001). Similarly, PolgA females were found to be 17% lighter than WT females (p<0.05) (Figure 1A). In terms of slowness, PolgA males exhibited a 23% reduction in walking speed compared to WT males (p<0.001), while PolgA females showed a 14% decrease compared to WT females (p<0.05) (Figure 1B). Although there was no significant difference in grip strength among males, PolgA males exhibited an 11% decrease in grip strength, whereas there was a 23% decrease in grip strength among females compared to WT females (p<0.05) (Figure 1C). In the case of physical activity, there was an 11% and 18% decrease for males and females respectively (p<0.05) between PolgA and WT mice (Figure 1D). Hence, the age-related differences were notably prominent in PolgA mice, with the differences being more pronounced in females around 40 weeks, especially in terms of mobility (walking speed and distance) and grip strength.

**Figure 1:**
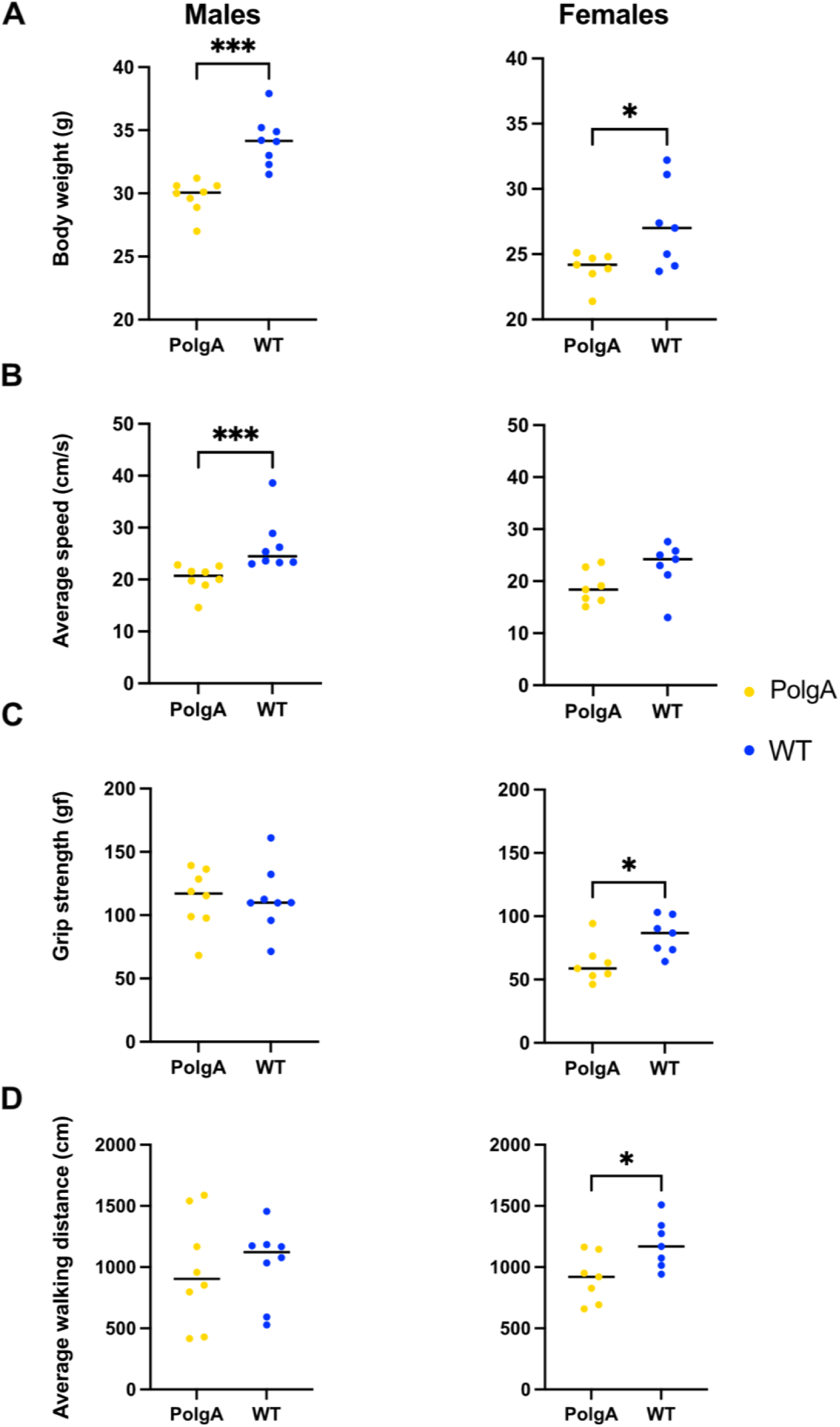
Changes in the musculoskeletal health parameters in PolgA mice: Comparison of each criteria: A) body weight B) average walking speed C), max grip strength and D) average walking distance used to assess frailty phenotype between PolgA mice and WT littermates at 40 weeks. Data points represent individual mice, with the horizontal line indicating mean, (n=7-8 mice/group). Significance was determined by the two-tailed Student’s t-test or Mann-Whitney test, (*p<0.05, **p<0.01, ***p<0.001).

To assess the frailty of PolgA mice, cutoff values, and the ranking order of mice for each criterion from best to worst, were identified (Figure 2A-B). Mice were considered frail if their performance fell below the cutoff values for the designated parameters for body weight (31.3 g, 24.4 g), grip strength (101.4 gf, 73.8 gf), average speed (23.3 cm/s; 21.6 cm/s) and average distance (769.1 cm, 1025.9 cm) for male and female mice respectively using the reference values obtained from WT mice around 40 weeks. According to these criteria, 5 PolgA male mice and 4 PolgA female mice were categorized as frail, while 3 PolgA male and female mice were classified as pre-frail. None of the PolgA mice were identified as non-frail. In contrast, among the WT mice, 2 males and females were designated as pre-frail, and 6 males and females were categorized as non-frail based on the same criteria. Regardless of sex, none of the WT mice were identified as frail (Figure 2A-B). In addition, these cut-off values obtained from WT mice for each sex were utilized as a reference to assess the prevalence of frailty in PolgA mice (Figure 2C). 62.5% of PolgA male mice were classified as frail (5 out of 8 mice), while 37.5% were categorized as pre-frail (3 out of 8 mice). On the other hand, 57.1% of PolgA females were identified as frail, while 42.9% were designated as pre-frail. In contrast, the majority of the WT males, 85.7%, were designated as non-frail while only 14.3% of them were identified as pre-frail. Similarly,75 % of WT females were categorized as non-frail and 25% as pre-frail (Figure 2C). The difference in frailty category (frail, pre-frail, and non-frail) distribution between PolgA and WT mice is statistically significant by the Fischer-Exact test for both females (p=0.0014) and males (p=0.0031), indicating the notable effects of genetic or physiological differences of these mice on their frailty status rather than random variability (Figure 2C).

**Figure 2:**
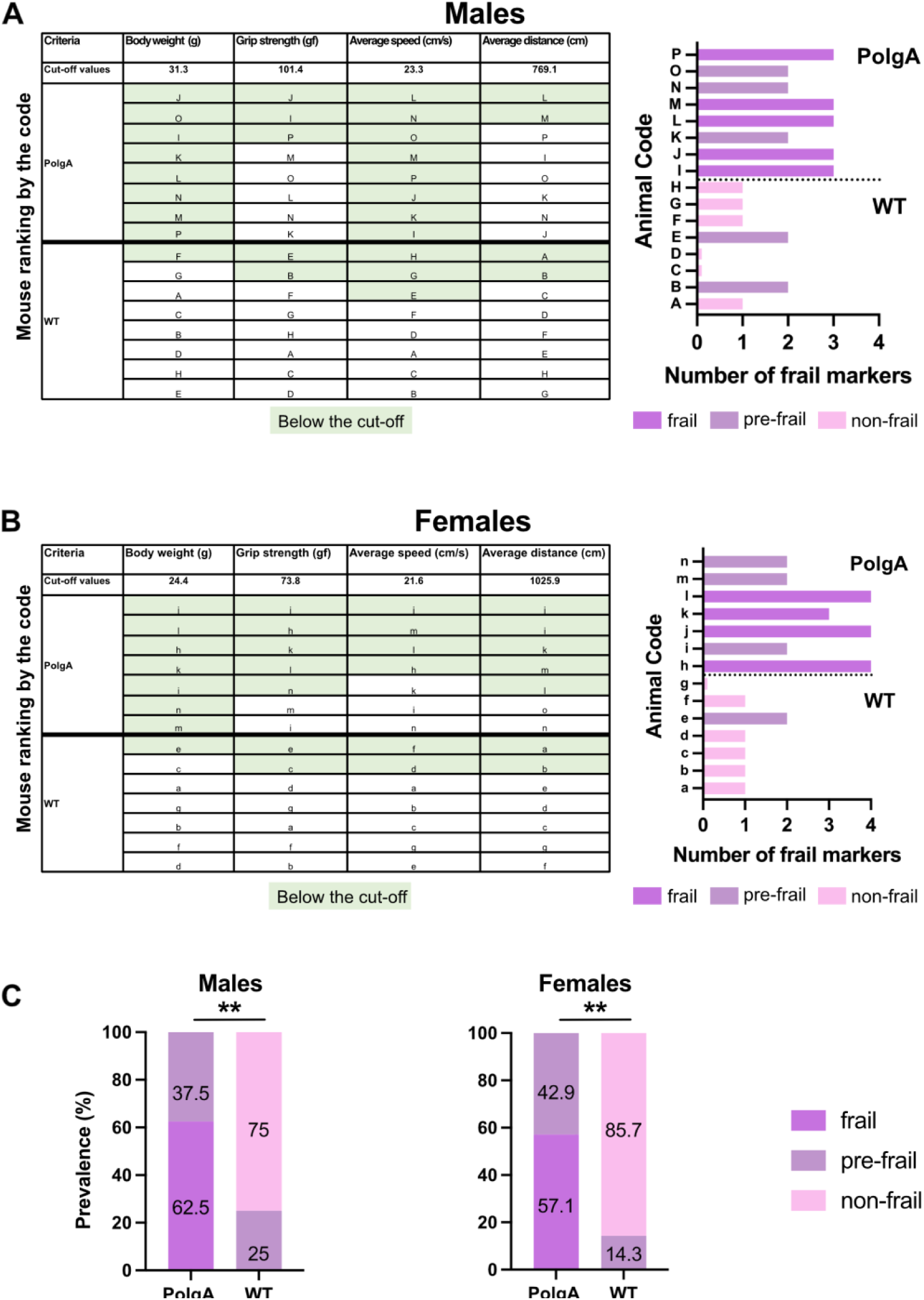
Frailty status of PolgA mice at 40 weeks. A) The mice were coded A-P for males and a-o for females and ranked by their performance. Frailty for PolgA mice of both sexes was calculated using reference values obtained from WT mice for each criterion. The green highlight indicates the mice in the below 20% of the performance for each criterion except body weight (below 10%). B) The number of frailty markers for each mouse at 40 weeks was presented. Frailty was defined if a mouse exhibited three or more of the criterion markers below the cut-off values. Pre-frailty was assigned if a mouse displayed two frailty markers, while non-frailty was indicated if a mouse had one or no markers. C) Prevalence for frailty was determined using the cutoff values obtained from 40-week-old WT mice for each sex and criterion categorized as frail, pre-frail, and non-frail. Data represented as percentages, (n=7-8 mice/group). Significance was determined by Fischer-exact test (*p<0.05, **p<0.01, ***p<0.001).

To identify the morphological differences between the mice identified as frail, pre-frail, and non-frail, we systematically compared these subgroups by evaluating each criterion to assess frailty. No distinct difference was observed in the body weight of PolgA and WT mice across frailty categories except pre-frail WT mice (2 data points) which weighed heavier than pre-frail PolgA mice (3 data points). However, non-frail female WT mice exhibited a higher body weight compared to frail and pre-frail mice. Similarly, pre-frail PolgA females weighed more than frail females, although the differences were not statistically significant (Figure 3A). Regarding slowness, there were no statistically significant distinctive variations in walking speed among the subgroups of PolgA and WT males and females. However, WT pre-frail male mice (2 data points) walked significantly faster than their pre-frail PolgA counterparts (3 data points). In contrast, female PolgA frail mice indicated a trend for slower walking speed in comparison to the pre-frail group while no variance in walking speed among WT females was observed (Figure 3B). For comparison of the weakness, pre-frail PolgA mice manifested a tendency towards higher grip strength compared to frail PolgA mice, regardless of sex. Whereas non-frail WT mice exhibited a trend for higher grip strength compared to pre-frail mice for both sexes (Figure 3C). A similar pattern was observed for comparison of physical activity assessed by average walking distance. Pre-frail PolgA mice indicated a pattern for increased walking distances compared to frail PolgA mice, irrespective of sex for both groups while non-frail WT mice covered longer distances compared to their pre-frail counterparts (Figure 3D). The results revealed subtle yet distinct morphological differences across the subcategories (non-frail, pre-frail, and frail) of WT and PolgA mice in evaluated parameters such as body weight, grip strength, and walking speeds. These differences were observed even though these groups were similarly categorized by frailty levels.

**Figure 3:**
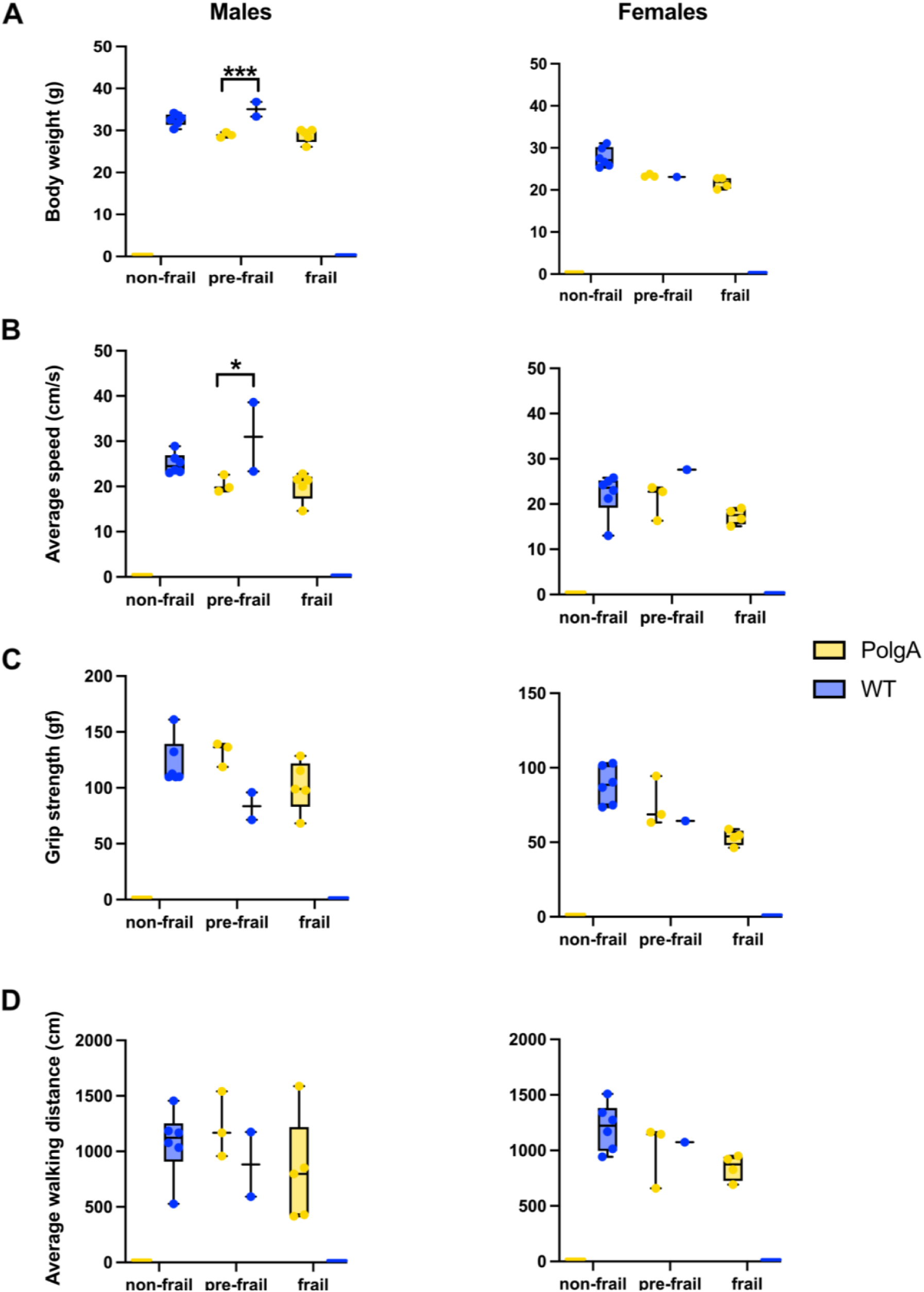
Morphological differences between frail, pre-frail, and non-frail mice. PolgA and WT mice that were identified as frail, pre-frail, and non-frail at 40 weeks were compared for each criterion used to evaluate the frailty phenotype: A) body weight, B) average speed, C) grip strength, D) average walking distance. Data represented as mean differences, (n=2-6 mice/group). Significance was determined by individual comparisons by the Two-stage linear step-up procedure Benjamini, Krieger, and Yekutieli (*p<0.05, **p<0.01, ***p<0.001).(Benjamini et al., 2006)

Furthermore, individual frailty scores were calculated as percentages, determined by dividing the total number of failed tests by the total number of tests performed for each group (Martinez de Toda et al., 2018) (Supplementary Table 1). PolgA males had a frailty score of 65.6%, while females had a score of 75%. In contrast, the frailty scores for WT mice were 25% for both males and females, indicating a higher level of frailty in female PolgA mice compared to males and WT females.

The calculation of frailty scores has revealed that the key contributors to frailty in our study include alterations in body weight, walking speed, and grip strength (Supplementary Table 2). Therefore, we compared these primary contributors used to evaluate frailty across different ages (Figure 4). Body weight gradually increases with aging in PolgA and WT mice from 12 to 40 weeks, with a more pronounced increase in WT mice. However, PolgA mice begin to experience weight loss after 40 weeks for both sexes. The differences in body weight between PolgA and WT mice become statistically significant at 34 (p<0.0001) and 40 weeks for males (p<0.001), while for females this significance is observed only at 40 weeks (p<0.01) (Figure 4A). Considering the slowness, there was no distinct difference between PolgA and WT mice males until 40 weeks. However, at 40 weeks, a significant decrease in walking speed is observed between WT and PolgA mice (p<0.01). In females, although there is a trend of decreasing speed with age in PolgA females, there is no notable difference between WT mice at any age (Figure 4A). Regarding the grip strength, there were no notable differences in males. However, in females, PolgA mice showed a gradual decrease over time, and at 40 weeks, this decrease was significant compared to WT females (p<0.05) (Figure 4A).

**Figure 4:**
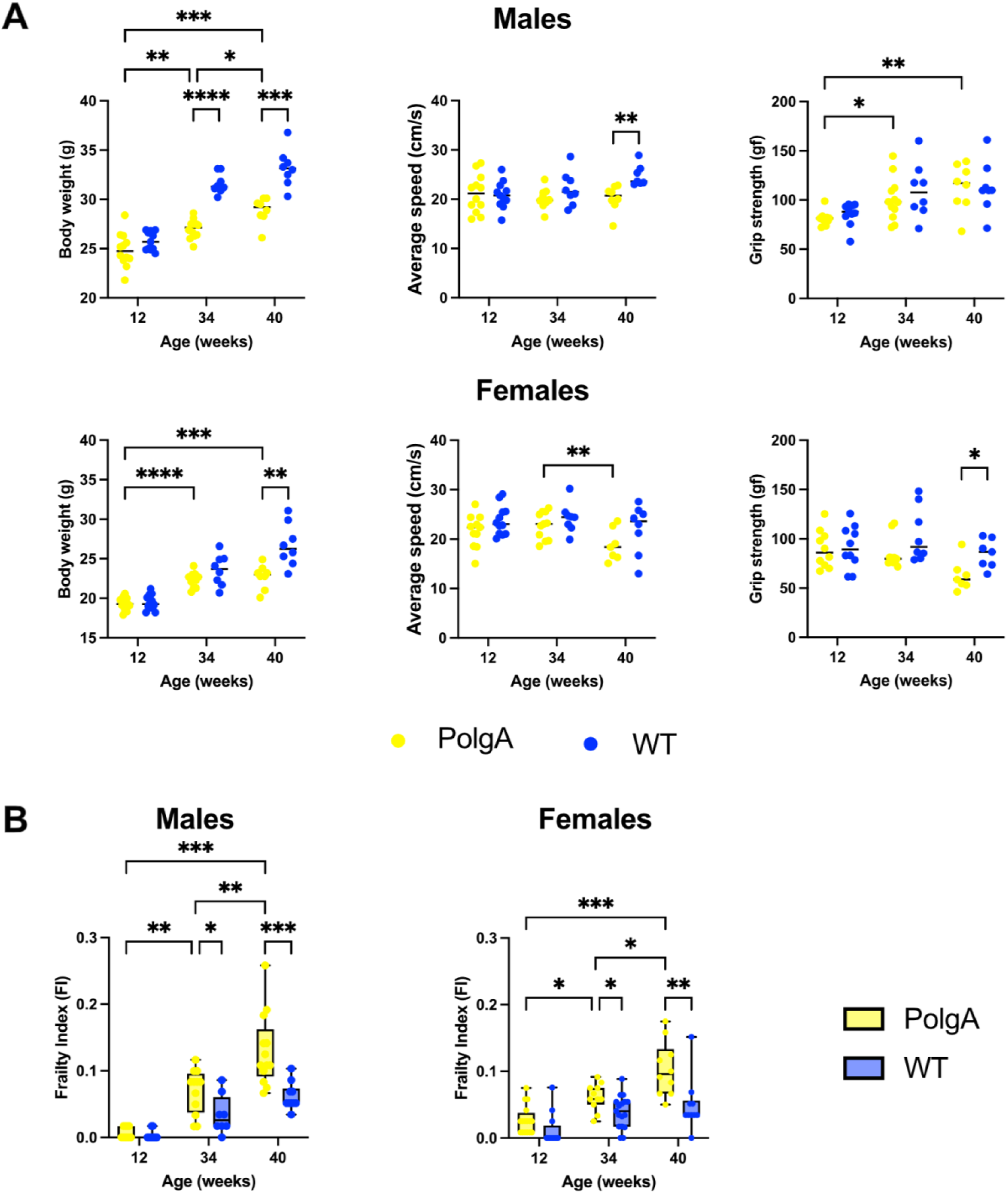
Comparison of key frailty phenotype parameters and frailty index across various age groups in PolgA mice. A) Assessment of the primary contributors used to evaluate frailty: body weight, slowness (average speed), and weakness (grip strength) B) Frailty index (FI) in PolgA and wild type (WT) at 12, 34, and 40 weeks. Data points represent individual mice, with the horizontal line indicating mean; box plots display max and min values, (*n* = 7-10 mice/group). Significance was determined by two-way ANOVA with Bonferroni or Sidak’s post-hoc tests (*p<0.05, **p<0.01, ***p<0.001).

Finally, we assessed FI in PolgA mice at these various ages. Our results showed a trend that the frailty of PolgA male and female mice increased with age. PolgA male mice demonstrated a higher FI compared to their WT (p<0.001) and female counterparts (p<0.01) at 40 weeks while no significant differences between the genotypes were observed at earlier time points (Figure 4B). This particular outcome might be associated with females exhibiting greater impairments in physical functions at 40 weeks while males accumulating a higher cumulative burden of health deficiencies although they performed better in these physical function measures.

## Discussion

Frailty measurement, utilizing both the FP and FI, is crucial for advancing our understanding of the aging process and developing interventions that can enhance health span. The FP is typically defined by physical characteristics such as reduced activity and weight loss, while FI, indicates a broad range of health deficits, and offers a comprehensive (Rockwood et al., 2005, Fried et al., 2001). Both approaches were employed to assess vulnerability to adverse outcomes; however, the Frailty Index (FI) has been suggested to be more precise in predicting mortality and physical performance in humans (Rockwood et al., 2007). Previous research on frailty in mice underscores the reliability of these measurements in capturing comparable aging phenomena observed in humans, demonstrating their critical role in translational aging research (Baumann et al., 2020, Liu et al., 2020, Parks et al., 2012). Furthermore, Feridooni et al. particularly investigated the reproducibility of FI scores done by different raters and concluded that mice have overall comparable FI scores (Feridooni et al., 2015). Consequently, it is clear that the two measurements are not equivalent. We opted for FP assessments but also conducted FI assessments for PolgA mice. As previously noted, the FI lacks the ability to fully capture the functional capabilities of the mice, yet it has provided indications of aging. We chose to maintain both measurements to illustrate how these mice not only exhibit signs of aging but also differ physically based on genotype and sex.

In this study, we applied the human frailty phenotype approach (Fried et al., 2001) to evaluate frailty characteristics of prematurely aging PolgA mice by assessing body weight loss, impaired strength, reduced walking speed, and limited physical activity. By applying this approach, we were able to identify the onset of frailty in PolgA mice and further elucidate the sex differences. PolgA mice exhibited a more pronounced physical decline, characterized by reduced body weight, average speed, grip strength, and walking distance. These parameters, used to assess frailty were notably more evident in females. Moreover there

Previous studies have shown that frailty in WT mice typically begins at 17 months, with onset often observed at around 27 months (Baumann et al., 2020). In contrast, PolgA mice exhibit frailty as early as 40 weeks, which roughly translates to 20-23 months in C57BL/6 mice; this timeline is analogous to humans developing frailty between 65-70 years of age (Fried et al., 2001, Dutta and Sengupta, 2016, Justice et al., 2016).

Moreover, by 29 months, 52.4% of WT mice were observed to be frail and another study reported that the frailty rates of 44.4%, for males and 73.7% for females at 26 months (Baumann et al., 2019, Baumann et al., 2020). Using the same assessment method, the prevalence of frailty was calculated for PolgA mice. In the PolgA males, which included 8 mice, 5 were identified as frail, resulting in a prevalence rate of 63%. In contrast, the analysis of the female cohort, consisting of 7 mice, revealed that 4 were frail, corresponding to a 57% prevalence rate. Although the prevalence rate is higher in PolgA males, females exhibit a more pronounced physical decline. In addition, we calculated frailty scores for PolgA mice resulting in 65.6 % for males and 75.0 % for females while the frailty score for the prematurely aging mice was previously reported as 40% (Martinez de Toda et al., 2018).

Collectively, our data show that PolgA mice have higher frailty scores compared to their age-matched WT counterparts irrespective of sex. Furthermore, the prevalence of frailty among PolgA mice at 40 weeks is comparable to that observed in WT mice at around 27 months of age. Considering the sex differences, female PolgA mice are frailer compared to male PolgA mice, with similar patterns observed in humans (Gordon et al., 2017), yet the limitations posed by a smaller sample size may impact the generalizability of this conclusion.

Moreover, there has been some controversy regarding increased frailty observed in female mice. While some studies have indicated no significant sex differences, others have demonstrated female C57BL/6 mice were frailer compared to males (Baumann et al., 2019, Whitehead et al., 2014, Parks et al., 2012). Furthermore, it was argued that the prevalence of frailty increases with age, ultimately resulting in early mortality among frail mice, regardless of sex (Baumann et al., 2019, Baumann et al., 2018, Kwak et al., 2020, Kane et al., 2018).

One limitation of our study is the small sample size contributing to non-significant results and altering the outcomes for the prevalence of frailty. In addition, the average walking distance of males might be impacted by their single housing conditions, potentially leading to no notable differences between the groups. Moreover, our frailty phenotype assessment is based on the 4 criteria adopted from the mouse frailty phenotype approach, which was derived from the human approach (Fried et al., 2001, Liu et al., 2014). However, the mouse FP approach lacked a weight factor and did not include an established endurance test. On that note, we could have incorporated an endurance test into our studies to provide a more comprehensive overview of the frailty phenotype status of PolgA mice. However, the traditional treadmill running test used to evaluate exhaustion in animals has faced significant scrutiny from an animal welfare point-of-view due to the stress it imposes on the animals. Furthermore, our lab has successfully conducted a longitudinal study on female PolgA mice and their WT littermates, investigating health span indicators such as the onset and progression of osteopenia and sarcopenia, as detailed in (Scheuren et al., 2020b). This study utilized *in vivo* micro-CT to monitor changes in bone microarchitecture from 20 to 40 weeks of age, capturing the transition from young to frail status. Unfortunately, the time required to breed and obtain male PolgA mice—exceeding a year—prevented similar longitudinal tracking for male mice within our study timeframe. Future studies should extend to include male mice to determine whether similar observations would be found or to explore potential differences in the progression of these conditions between sexes.

In conclusion, our study effectively characterized the frailty phenotype in prematurely aging PolgA mice. Frailty was consistently observed at 40 weeks, evidenced by signs such as loss of body weight, decreased walking speed, grip strength, and overall physical activity. FI scoring corroborated these findings and highlighted an age-associated increase in frailty with the onset observed at 40 weeks in PolgA mice. Furthermore, our study revealed sex differences, with females exhibiting more pronounced physical decline than males. Our findings align with published literature detailing similar characteristics observed in wild-type (WT) mice and humans with the frailty phenotype, emphasizing the utility of PolgA mice as a relevant model for studying aging and associated disorders.

## Methods

### Animal models

All animal procedures were carried out by the regulations set by the local authorities (license: ZH35/2019) and approved by the Verterinäramt des Kantons Zürich, Zurich, Switzerland. A colony of mice expressing an exonuclease-deficient version of the mitochondrial DNA polymerase γ (PolgAD257A, B6.129S7(Cg)-Polgtm1Prol/J, JAX stock 017341, The Jackson Laboratory) was bred and maintained at the ETH Phenomics Center. The mice were housed under standard conditions, featuring a 12:12 h light-dark cycle, with access to maintenance feed–vitamin fortified pelleted diet (3437 KLIBA NAFAG Kaiseraugst, Switzerland) and water ad libitum, and three to five animals per cage. The mice were bred as previously described (Scheuren et al., 2020a) and genotyped before the experiments (Transnetyx, Cordova, USA). Homozygous PolgA mice and age- and sex-matched WT littermates (controls) were used for all experiments.

### Grip strength

Grip strength was assessed using a force tension apparatus (Grip Strength Meter, model 47200, Ugo Basile) approximately at 40 weeks in female and male PolgA and WT mice as previously described (Scheuren et al., 2020a). The mice held the stationary bar with their forepaws and were then gently pulled horizontally by their tail until they released their grip. This procedure was repeated five times for each mouse, and the maximum force (gram-force) value was used for the analysis. All measurements were conducted by the same trained operator.

### Gait analysis

Gait data was collected from female and male PolgA and WT mice around 40 weeks of age using the Noldus CatWalk XT system. Run speed, stand, duty cycle, swing speed, maximum variation, and stride length were analyzed to assess the gait characteristics of the mice.

### Physical activity

The average distance was measured from the Phenomaster experiment and used to assess physical activity level. The PhenoMaster metabolic cage system was utilized to record various physiological and behavioral measurements including the measurement of drinking and feeding amounts, CO_2_ and O_2_ consumption, walking speed, and walking distance(Singh et al., 2024). For this manuscript, only the average walking distance from the PhenoMaster analysis was used.

### Quantification of frailty index

The FI in the study was quantified by implementing the Mouse Clinical Frailty Index consisting of a comprehensive evaluation of 31 non-invasive clinical parameters (Whitehead et al., 2014). Each parameter except body weight and body surface temperature received a score of 0 for absence, 0.5 for a mild deficit, and 1 for a severe deficit. Body weight and body surface temperature were scored relative to the standard deviations from a reference mean in young adult mice (12 weeks old). However, body surface temperature was excluded in the calculation of the FI due to a lack of appropriate tools for measurement.

### Frailty phenotype criteria

To establish a frailty phenotype, we identified 4 criteria that are indicative of frailty: body weight loss, grip strength, walking speed, and physical activity. These criteria were deliberately chosen due to their similarity to the clinical criteria for assessing frailty in humans (Fried et al., 2001). By incorporating these key indicators, we aimed to create a comprehensive and reliable frailty phenotype that aligns closely with established clinical mouse and human frailty standards (Table 1). For each parameter, a 20% cut-off value was chosen, except for body weight. In the case of body weight, which was measured prior to the experiments, a 10% cut-off value was applied. This choice was made to maintain alignment with established human criteria and to ensure consistency in assessments between mice and humans, as indicated in the previous publication (Fried et al., 2001, Hill, 2009, Hill et al., 2009, Koch et al., 2016, Leibel, 2008).

**Table 1:** The criteria for defining mouse frailty for PolgA mice based on the human frailty criteria.

| Human frailty criteria | Mouse frailty criteria | Cut off values |
| --- | --- | --- |
| Unintentional weight loss | Body weight change:<br>Body weight (g)<br>- Males: 31.3 g<br>- Females: 24.4 g | Lowest 10% |
| Weakness - Low grip strength | Weakness:<br>Grip strength (gf)<br>- Males: 101.4 gf<br>- Females: 73.8 gf | Lowest 20% |
| Slow walking speed | Slowness:<br>Average walking speed (cm/s)<br>- Males: 23.3 cm/s<br>- Females: 21.6 cm/s | Lowest 20% |
| Low physical activity level | Low physical activity:<br>Average walking distance<br>(Liang et al.)<br>- Males: 769.1 cm<br>- Females: 1025.9 cm | Lowest 20% |

The varying cut-off values are designed to closely mirror the original clinical assessments used in human studies, while also balancing sensitivity (the ability to correctly identify those who are frail) and specificity (the ability to correctly identify those who are not frail).A smaller percentage change in body weight may be more indicative of frailty and its associated outcomes compared to similar percentage changes in grip strength or physical activity. Moreover, measures such as body weight tend to show less variability over time, unlike grip strength or walking speed, which can be affected by immediate factors like short-term illness, fatigue, or the time of day. Furthermore, our analysis, which used the 10% percentile as a cutoff value for body weight changes, revealed no significant differences when compared to the 20% cutoff values (Supplementary Table 3).

### Statistical analysis

Data analysis was performed with GraphPad Prism software. Fischer exact test to analyze the frailty category data where sample sizes were small. For controlling the false discovery rate in multiple testing of the frailty phenotype parameter within sub-frailty groups, we applied the Two-stage linear step-up procedure of Benjamini, Krieger, and Yekutieli, accounting for both fixed and random effects in the data. Continuous variables were assessed using the two-tailed Student t-test or the Mann-Whitney test when comparing two independent samples, and two-way ANOVA with Bonferroni or Sidak’s posthoc tests was utilized for comparing more than two groups. Statical significance was determined at p<0.05.

### Data availability

The authors declare that all data supporting the findings of the study are available within the paper and could be shared upon reasonable request from the corresponding author.

## Acknowledgments

We express our gratitude to ETH Phenomics Centre (EPIC) at ETH Zurich for their support and to Susanne Friedrich, the Head of Phenotyping module at EPIC, ETH Zurich, for her assistance in conducting grip strength and PhenoMaster experiments. We acknowledge Abhay Khosla and Shekinah Arpon for their help with the analysis of gait data. We also acknowledge ChatGPT, an AI language model developed by Open AI to provide guidance to improve the readability of this manuscript. After using this tool, the authors reviewed and edited the content as needed and took full responsibility for the content of the publication.

## Author Contributions

DY prepared figures, performed the data analysis, and drafted the manuscript. AS contributed to obtaining physical activity data while NM oversaw the analysis of the gait data. GAK designed and performed the phenotyping experiments used in the study for this manuscript. DY, NM, AS, EW, GAK, and RM, edited and revised the manuscript; DY, NM, AS, EW, GAK, and RM approved the final version.

