## Supplementary Files for "Unveiling Frailty: Comprehensive and Sex-Specific Characterization in Prematurely Aging PolgA Mice"

**Supplementary Table 1: Calculation of frailty score in PolgA mice and WT littermates:**

The frailty score was calculated using reference values obtained from WT mice for each sex and criterion with A representing the total number of mice below designated the cut-off value for the specific criterion, and B representing the total number of mice included in the analysis for that criterion. Frailty score is expressed as a percentage.

| **Criteria** | **Males** | | | | **Females** | | | |
|---|---|---|---|---|---|---|---|---|
| | **PolgA** | | **WT** | | **PolgA** | | **WT** | |
| | A | B | A | B | A | B | A | B |
| Body weight (g) | 8 | 8 | 1 | 8 | 7 | 7 | 1 | 7 |
| Average walking speed (cm/s) | 8 | 8 | 3 | 8 | 4 | 7 | 2 | 7 |
| Grip strength (gf) | 3 | 8 | 2 | 8 | 5 | 7 | 2 | 7 |
| Average walking distance (cm) | 2 | 8 | 2 | 8 | 5 | 7 | 2 | 7 |
| Total | 21 | 32 | 8 | 32 | 21 | 28 | 7 | 28 |
| Frailty Score (%) (A/B) x 100 | 65.6 | | 25 | | 75 | | 25 | |

**Supplementary Table 2: Calculation of frailty score in PolgA mice and WT littermates using 3 frailty criteria:**

The calculated frailty scores (abcd) for PolgA mice and WT littermates, derived from the method presented in Table 2, were extended to assess frailty using a combination of three criteria: Body weight (a), Average speed (b), Grip strength (c), and Average walking distance (d), representing a subset of the four criteria used in Supplementary Table 1.

| Criteria | Frailty Score (%) | | | |
|---|---|---|---|---|
| | Males | | Females | |
| | PolgA | WT | PolgA | WT |
| abcd | 65.6 | 25 | 75 | 25.0 |
| abc | 79.2 | 25 | 76.2 | 23.8 |
| abd | 75.0 | 25 | 76.2 | 23.8 |
| acd | 54.2 | 20.8 | 81 | 23.8 |
| bcd | 54.2 | 29.2 | 66.7 | 28.6 |

**Supplementary Table 3: Comparison of 10% vs 20% percentile cutoffs of body weight**

The table layout allows for a comparison of two cutoff values of body weight and their impact on the frailty status of WT mice. The frailty status of both male and female PolgA mice was unaffected by changes in the body weight cutoffs; in both scenarios, all mice were classified as frail. A statistical comparison was conducted using the Wilcoxon matched-pairs signed rank test, which indicated no significant difference in frailty status between mice with different cutoff values.

| **WT Males** | | | | | |
|---|---|---|---|---|---|
| **Cutoff percentile** | **Body weight (g)** | **frail mice** | **pre-frail mice** | **non-frail mice** | **p-value** |
| 10% | 31.3 | 0 | 2 | 6 | 0.5 |
| 20% | 32.0 | 0 | 3 | 5 | |
| **WT Females** | | | | | |
| **Cutoff percentile** | **Body weight (g)** | **frail mice** | **pre-frail mice** | **non-frail mice** | **p-value** |
| 10% | 24.6 | 0 | 1 | 6 | >0.9999 |
| 20% | 25.5 | 0 | 2 | 5 | |
